# Light-weight Electrophysiology Hardware and Software Platform for Cloud-Based Neural Recording Experiments

**DOI:** 10.1101/2021.05.18.444685

**Authors:** Kateryna Voitiuk, Jinghui Geng, Matthew G. Keefe, David F. Parks, Sebastian E. Sanso, Nico Hawthorne, Daniel B. Freeman, Mohammed A. Mostajo-Radji, Tomasz J. Nowakowski, Sofie R. Salama, Mircea Teodorescu, David Haussler

## Abstract

**Objective:** Neural activity represents a functional readout of neurons that is increasingly important to monitor in a wide range of experiments. Extracellular recordings have emerged as a powerful technique for measuring neural activity because these methods do not lead to the destruction or degradation of the cells being measured. Current approaches to electrophysiology have a low throughput of experiments due to manual supervision and expensive equipment. This bottleneck limits broader inferences that can be achieved with numerous long-term recorded samples.

**Approach:** We developed Piphys, an inexpensive open source neurophysiological recording platform that consists of both hardware and software. It is easily accessed and controlled via a standard web interface through Internet of Things (IoT) protocols.

**Main Results:** We used a Raspberry Pi as the primary processing device and Intan bioamplifier. We designed a hardware expansion circuit board and software to enable voltage sampling and user interaction. This standalone system was validated with primary human neurons, showing reliability in collecting real-time neural activity.

**Significance:** The hardware modules and cloud software allow for remote control of neural recording experiments as well as horizontal scalability, enabling long-term observations of development, organization, and neural activity at scale.

## 1. Introduction

Extracellular voltage recordings from *in vitro* cell cultures support the investigation of neural activity and dynamics. These recordings allow us to assess information processing in complex neuronal networks and enable discovery on a scale from single neuron firing patterns to local and long-range functional connectivity, network synchrony, and oscillatory activity [1–6].

Longitudinal recordings are essential to capture features of neurodevelopment and dynamics: basic physiological properties of neuron development, how 2D and 3D cultures grow and change activity patterns, and what rhythms the activity may follow [7–10]. Recordings across time are essential to study response to electrical or drug stimulus over weeks and months. Wider application are used for drug discovery and genetic screens.

The further combination of longitudinal recordings and large numbers of parallel experimental replicates allow investigations to progress significantly faster and makes new experiments feasible [11]. Also, scaling up experiments generates the large volume of data necessary for taking advantage of Machine Learning algorithms and creates a faster turnaround between hypothesis, experiment, and re-testing [12]. *In vitro* culture models serve as a flexible system that are much easier to scale up than animal models, especially when paired with developments in robotic automation, microfluidics, and probes [13–16].

Longitudinal recordings from multi-channel experiments demand vast amounts of data and memory. The data is challenging to manage, especially since out-of-the-box hardware and software are often offline. Storage on physical disks usually requires manual monitoring to prevent running out of disk space and laborious transfer of data for backup or processing. Furthermore, many recording systems require a designated workspace for experiments with a physical computer nearby with cables or wireless transmission to stream data. Several open-source efforts have been created to provide more affordable and modifiable recording equipment [17–21]. However, no software solutions exist to easily manage and control a large amount of electrophysiology equipments and data at once.

Recent advances in commodity hardware allow for more affordable computing devices. The Internet of Things allows many devices to come online when needed and be relinquished when not needed, and protocols have been developed to effectively and securely manage and communicate with these devices. Affordable, internet-connected devices have already been developed for ECG, EEG, EMG, and heart rate variability monitoring [22–27]. Furthermore, commodity cloud compute from major companies as well as academic coalitions [28] has become widely available and many tools for downstream analysis to process voltage recordings are already offered online [14, 29–32]. However, data acquisition for *in vitro* cultures remains relatively isolated, as no platform exists to stream data online to link with these analysis infrastructures. One solution is to write software add-ons for existing data acquisition systems. However, not all existing data acquisition systems are flexible or open in terms of data formats, programmability, and remote control. Additionally, channel count and price range are not always suitable for the desired application.

To address these issues we created Piphys, an all-in-one electrophysiology and processing system that can simultaneously record data from multiple channels in the *μ*V scale and stream it to the cloud. The user interacts with the device through a dashboard website to view data and control experiment parameters. Both hardware and software are made available as open source.

Piphys is based on a Raspberry Pi computer and eliminates the need for a desktop or laptop computer to manage an electrophysiology experiment or for an operator to be present in the lab to start a recording. The Raspberry Pi comes with a Unix-based operating system that can be easily programmed with many existing software libraries and tools. Overall, the low price and extreme flexibility of the Raspberry Pi significantly lowers the cost of the entire electrophysiology system, providing an opportunity for broader education and research opportunities.

Piphys can be used with a wide range of electrode probes including, but not limited to, rigid 2D and flexible 3D microelectrode arrays (MEAs) [33], silicon probes [34], and tetrodes. The system is built for long-term experiments with the goal of full automation using programs that can optimize experimental variables. Here we detail the Piphys system’s functionality and validate its accuracy and reliability for measuring neural activity.

## 2. Piphys System Design

The Piphys hardware records from neural tissue remotely using our versatile circuit board connecting to Intan RHD series recording chips to perform highly sensitive analog to digital conversion. Data from the Intan can be optionally preprocessed on-site using a Raspberry Pi computer and streamed to a cloud service where deeper sorting and analysis of detected spikes can be performed. Spike sorting analysis measures neural activity changes over time in individual neurons and networks of neurons, using features like spike waveform, frequency of activity, and correlation to the activity of nearby neurons.

### Hardware design

#### Design elements (choice of platform)

The key physical innovation in Piphys is a hardware expansion board that enables a Raspberry Pi computer to interface with an Intan RHD2132 bioamplifier chip to perform electrophysiology.

The Raspberry Pi Model 3 B+ is a low-cost, small-scale, single-board computer. It has a quad-core ARM Cortex-A53 processor with an Input/Output system. It can be programmed to interface with customized hardware with a standard data communication protocol. It also has an expandable memory space configured by a removable SD card.

The Intan RHD2132 bioamplifier chip is the key driver of the shield biopotential-sensing functionality. The chip amplifies voltage signals sensed by the electrodes and converts the analog signals to digital values for storage inside the Raspberry Pi computer.

#### Circuit design

An expansion shield connects the Raspberry Pi to the Intan RHD 32-channel recording headstage containing the Intan RHD2132 bioamplifier chip. The chip is configured to use low-voltage differential signaling (LVDS) to reduce the effects of noise and electromagnetic interference (EMI) and allow increased cable length. However, the Raspberry Pi communicates using complementary metal-oxide-semiconductor (CMOS) level logic. To translate between the two signal types, the expansion shield uses the SN65LVDT41 chip from Texas Instruments. The SN65LVDT41 chip has four LVDS line drivers and one LVDS line receiver to control data lines required to communicate with the Intan chip over its Serial Peripheral Interface (SPI).

Besides translation between signal types, the expansion shield provides different levels of power derivative from the +5V source input. The +5V input powers both the Pi and shield, and can be supplied either through the power barrel on the shield or through the micro-USB on the Pi for flexibility. On the shield, the power source is filtered through ferrite beads to remove high-frequency power line noise. The +5V source is converted to a +3.5V source for the Intan RHD2132 bioamplifier chip and a +3.3V for the SN65LVDT41 chip. Conversion is performed by low-noise linear voltage regulators to smooth and isolate any fluctuations from the power supply.

#### Connection to electrodes

Electrodes are connected to the Intan RHD 32-channel recording headstage. For experiments reported here, we created a connection to a commercially available 6-well multi-electrode array (MEA) plate from Axion Biosystems. However, any other electrode system fitting an Omnetics 32-pin connector is compatible. The design can be adapted to custom and commercial MEAs of different form factors using adapter boards.

The Axion electrode plate mates its bottom contacts to spring finger pins on our designed adapter board. The parts are aligned using a custom holder consisting of a plastic interior surrounded by aluminum plates and compressed together by screws on four corners. The plastic holder has a slot to hold the adapter board and a groove to align the plate in the correct position. The aluminum plate casing prevents warping of the plastic and ensures even pressure compressing the plate and connector on both sides. The compressing holder provides consistent mating of spring finger pins to electrode contacts on the plate.

### Software design

The Piphys system runs custom software to perform: (1) communication with the Intan RHD2132 bioamplifier chip, (2) buffering and file storage of recorded voltage data locally, (3) real-time data streaming and plotting on the online dashboard, and (4) experiment control from the dashboard. In order to stream data, interact with data being recorded, and control the device, we deployed Redis, Amazon IoT, and S3 as described in Methods.

To perform an electrophysiology recording, the user can configure the sampling rate and start the experiment from the dashboard. Once started, the neural cell activity is firstly digitized and sampled by the Intan RHD2132 bioamplifier chip in 32 channels. Raspberry Pi stores the data on local memory and also streams it to Redis for real-time visualization on the online dashboard. For data integrity and upload efficiency, raw data is saved every 5 minutes on local memory and streamed every 10 seconds to Redis. Once the recording ends, all local data files are uploaded to S3 for permanent storage, and data is further backed up to Amazon Glacier for long-term archiving. Local data files on the Pi auto-erase every 14 days to release memory. To view a dated recording, the user can select and pull the data files from S3 to the dashboard for display (Figure 3).

**Figure 1.**
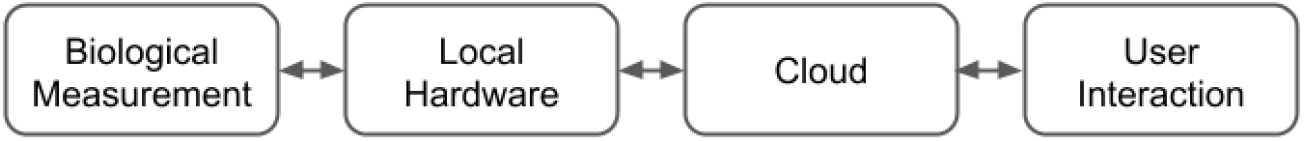
Cloud-based experiment paradigm: biological measurement and local hardware are presented to the user through the cloud, such that experiment management and control can be administrated remotely and may be automated by a computer program.

**Figure 2.**
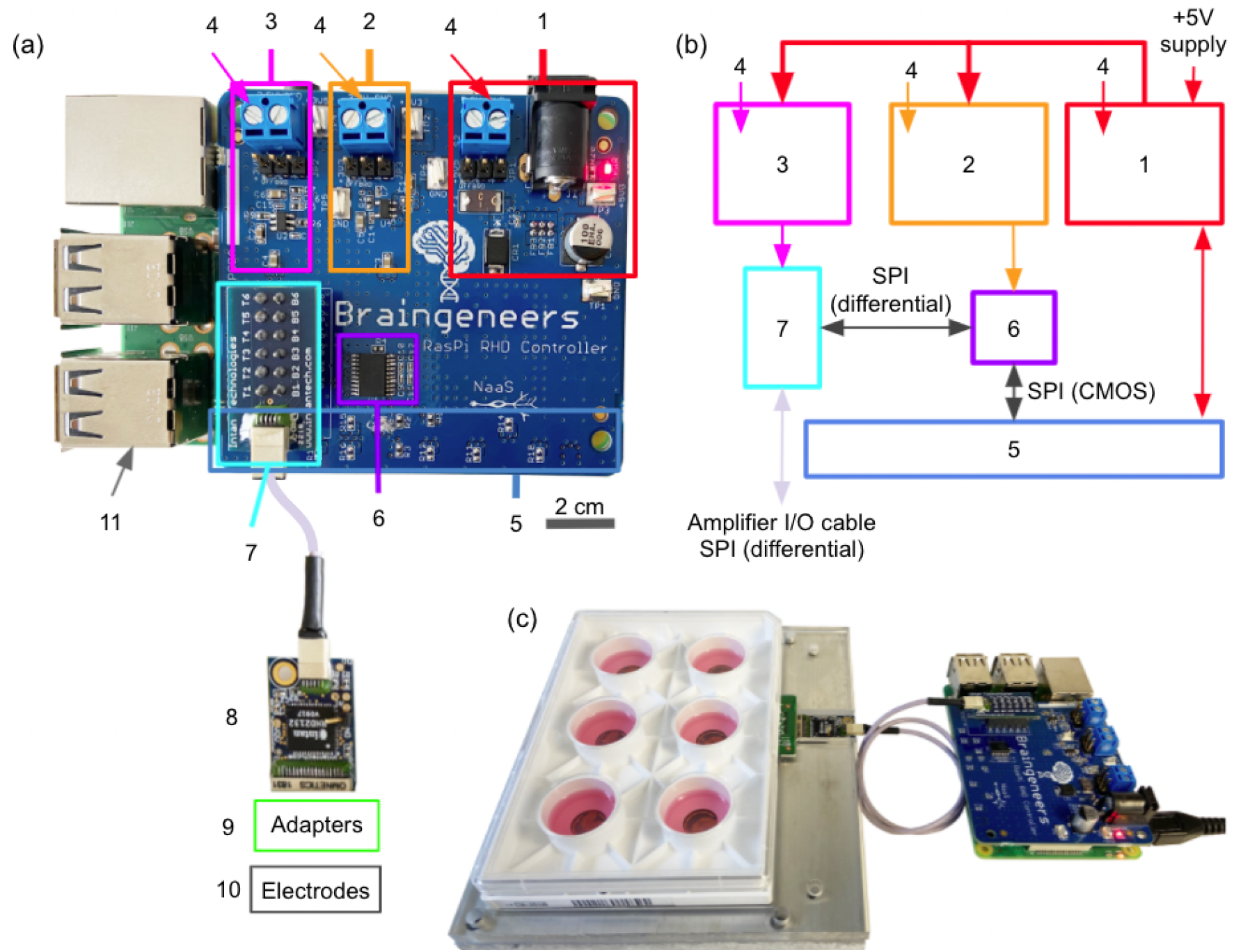
Piphys hardware components. **(a)** Expansion shield (blue board) attached on top of Raspberry Pi (green board). **(b)** Logic level connection. **(c)** Example interface with standard 6-well electrode plate. (1) +5V logic, (2) +3.3V logic, (3) +3.5V logic, (4) External supply inputs, (5) Raspberry Pi input/output pins (bottom), (6) LVDS converter, (7) Intan RHD adapter, (8) Intan RHD 32-channel recording headstage containing Intan RHD2132 bioamplifier chip, (9) Optional adapter board to electrodes, (10) Multiple electrode types possible, (11) Raspberry Pi computer (bottom).

**Figure 3.**
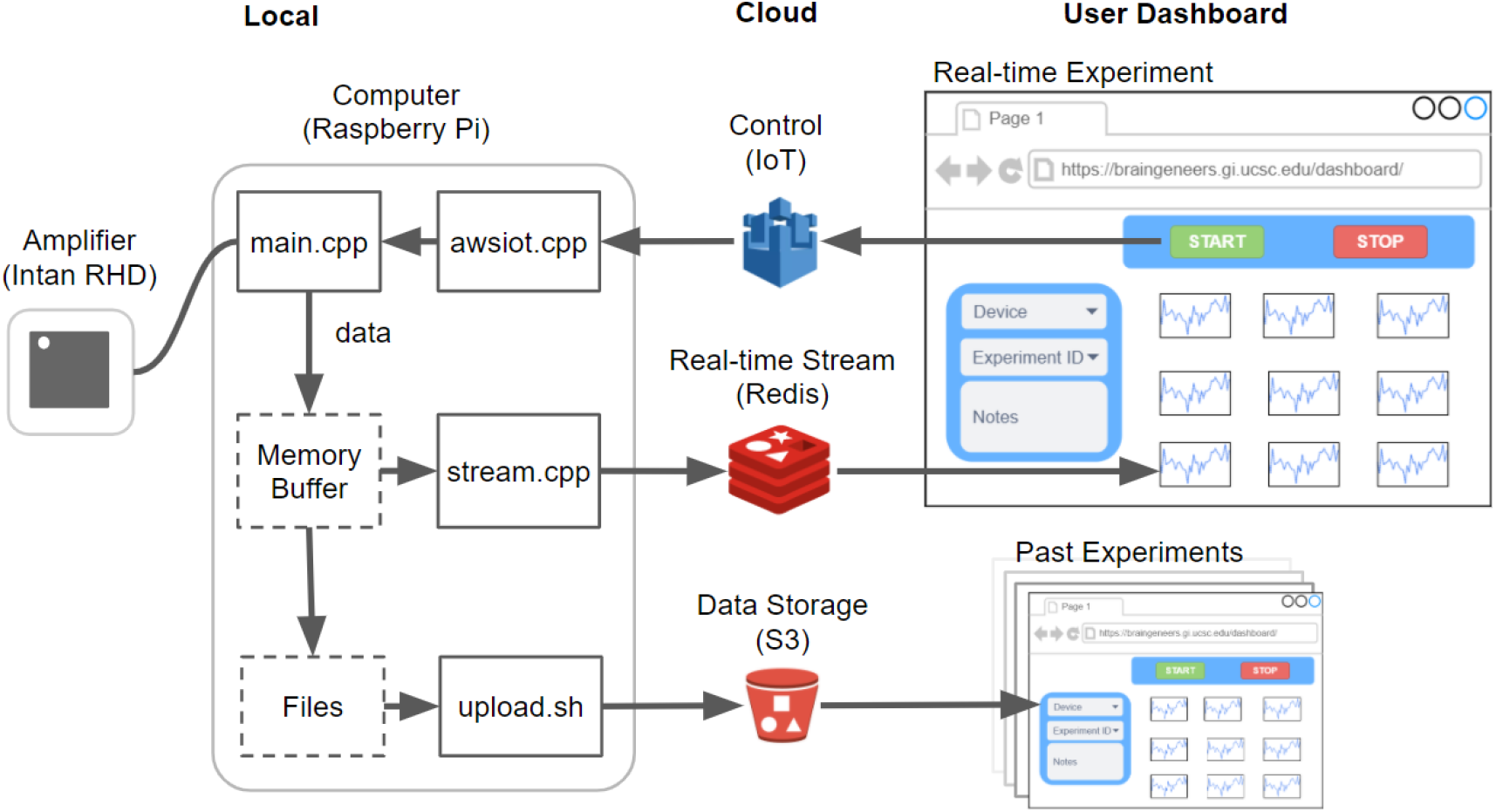
Software overview. The software that runs on the local Raspberry Pi device communicates with the Intan RHD2132 bioamplifier chip to stream and store the digitized neural signal. Concurrently, it pushes the signal to Redis for real-time visualization on the online dashboard. Datasets are also uploaded to S3 after each recording for permanent storage and access. Experimental control such as ‘start’, ‘stop’, and variable configuration is sent from the dashboard through Amazon IoT to the local device. Past experiment data can also be browsed using records from S3.

**Figure 4.**
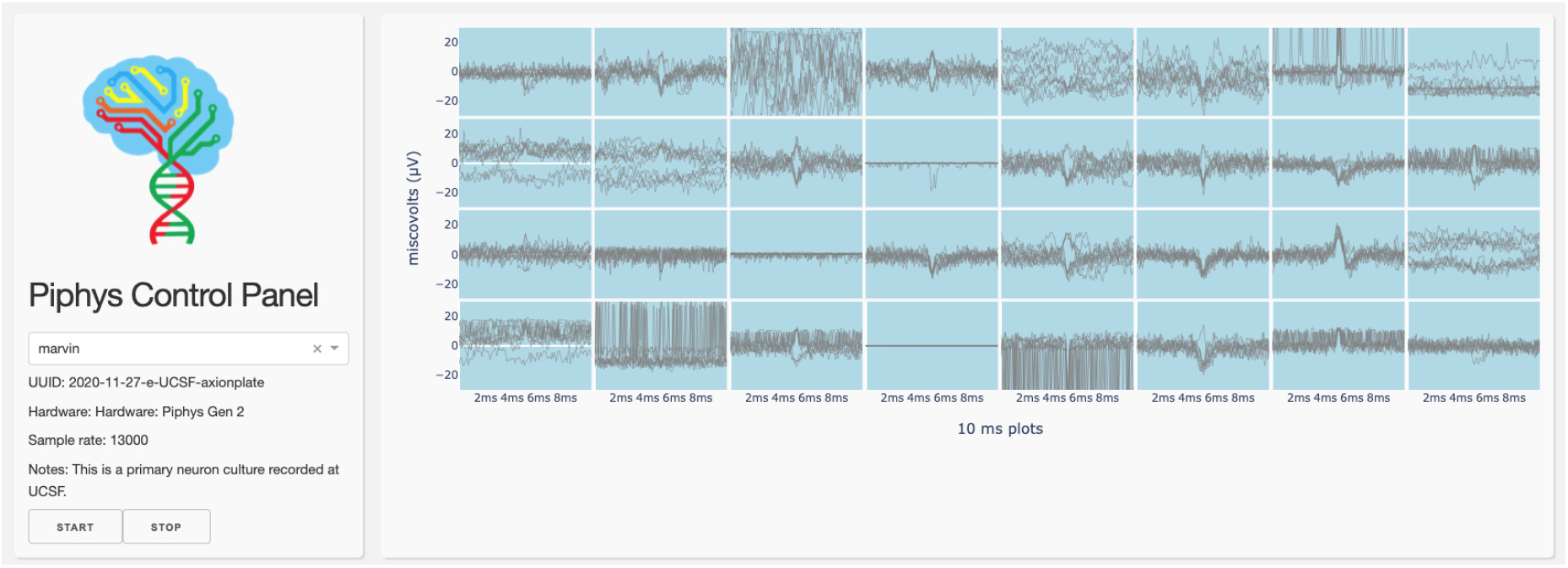
Dashboard. A control panel interface is displayed through the browser running spike detection by thresholding.

#### Communication with hardware

Communication between Raspberry Pi and Intan RHD2132 bioamplifier chip uses Serial Peripheral Interface (SPI). SPI is a fast and synchronous interface that is widely used in embedded systems for short-distance data streaming. It is a full-duplex master-slave-based interface where both master and slave can transmit data at the same time. The protocol for both Raspberry Pi and Intan RHD2132 bioamplifier chip is a four-wire interface: Clock (SCLK), Chip select 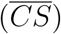, Master-Out-Slave-In (MOSI), and Master-In-Slave-Out (MISO). In Piphys, the Raspberry Pi acts as the master device and generates a clock signal and recording commands to configure the Intan RHD2132 bioamplifier chip through MOSI. The Intan chip responds as slave and sends the digitized data back by MISO. The chip allows configuration of sampling rate and bandwidth of the low-noise amplifiers. The 32 channels on the chip are sampled sequentially with available sampling rate options from 2 kHz to 15 kHz per channel. The amplifiers give 46 dB midband gain with lower bandwidth from 0.1 Hz to 500 Hz, and upper bandwidth from 100 Hz to 20 kHz.

#### Online dashboard

Users interact with Piphys devices through a web browser application, referred to as the Graphical User Interface (GUI). The GUI allows a user to initiate a recorded experiment and monitor electrical activity on each channel. Programatically, the GUI mimics an IoT device that sends messages to other devices (i.e., Piphys units) and listens to their corresponding data streams in a high-performance Redis database service. The Piphys device produces a single data stream to Redis, and many users can view the stream from the Redis server. Therefore, many users can monitor and interact with a particular Piphys device without additional overhead placed on that device.

Users can be located anywhere on the Internet without concern for where the physical Piphys device is or which network it is on. We routinely perform electrophysiology experiments from Santa Cruz on a Piphys-connected device that is located 90 miles away in San Francisco.

When a new user opens the browser GUI, the web application queries the AWS IoT service for online Piphys devices to populate a device dropdown list. When the user selects a device from the dropdown, an MQTT ‘ping’ message is sent to the relevant device every 30 seconds, indicating that a user is actively monitoring data from that device. As long as the Piphys device receives these pings, the Piphys device will continue to send its raw data stream to the central Redis service. When the Piphys device has not received any user messages for at least a minute, it will cease sending its raw data stream. This protocol ensures the proper decoupling of users from devices. The Piphys device is not dependent on a user gracefully shutting down.

While the Piphys device feeds raw data to the Redis service, data transformations are applied downstream by other IoT-connected processes. For example, the Piphys Control Panel displays a threshold spike sorted transformation of the raw data. This data transformation is an independent process that listens for MQTT requests for the raw data stream and transforms the raw stream into a stream containing the past ten spike events detected per channel. For channels with no detected spikes, a random sample of the channel is saved to the stream every 30 seconds to provide a sampling of the channel’s activity.

## 3. Detection of neuron activity

We tested the Piphys system for long-term recordings of human primary neurons. These neurons were cultured in an Axion CytoView MEA 6-well plate (Methods). After recording, the raw data was ingested to SpyKING CIRCUS software [31] for analysis. SpyKING CIRCUS is a semi-automatic spike sorting software that uses thresholding, clustering, and greedy template match approaches to detect single cell action potentials. Here, we show two types of results, first for single neuron recordings and second for a bursting neural network.

### Recording from primary neurons

After 14 days in culture in culture, primary neurons were recorded with the Piphys system and two commercially available systems: the Intan RHD USB interface board and the Axion Maestro Edge. After recording, all three datasets were filtered with bandpass filtering from 300 Hz to 6000 Hz and spike sorted with a threshold of ± 6 *μV*. Figure 5 shows a ten-second spike train from Piphys with dots highlighting detected spikes in the raw data.

**Figure 5.**
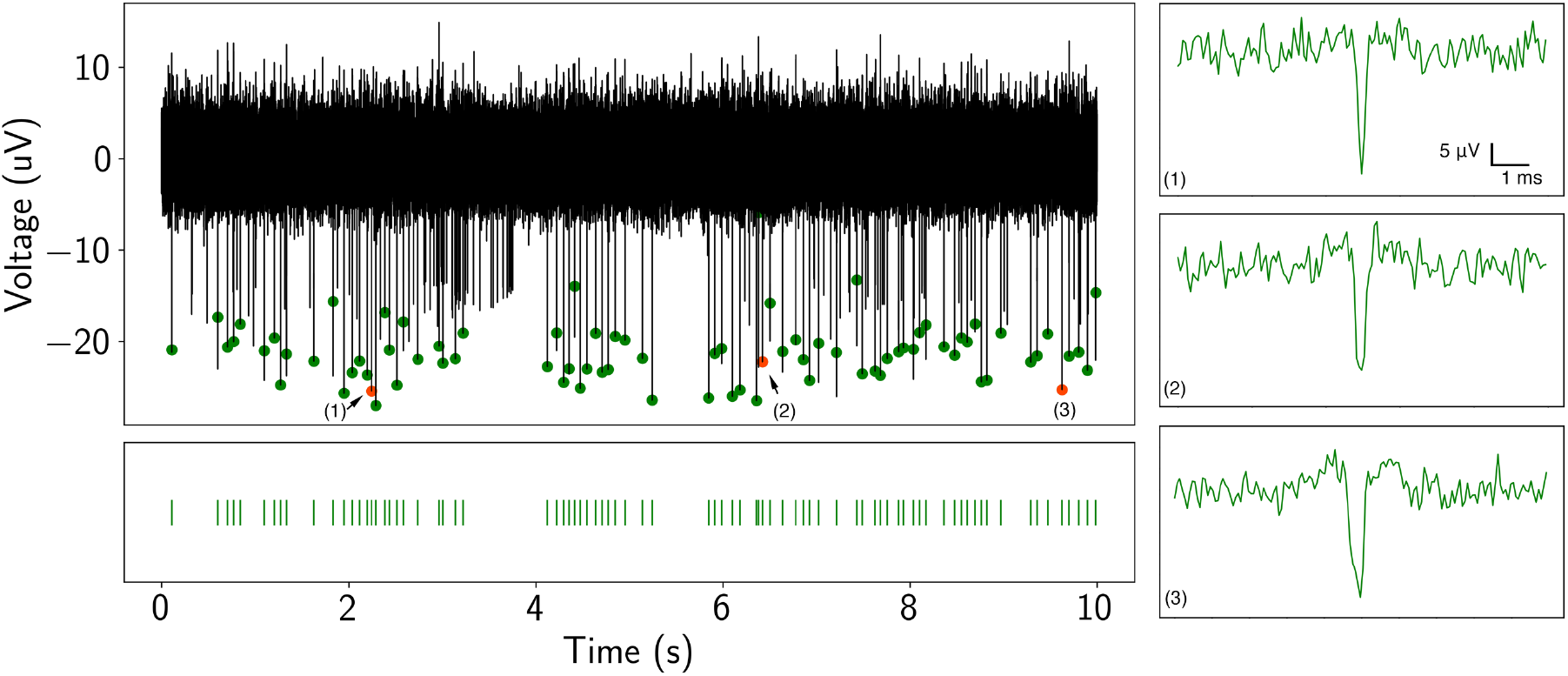
Detection of neuronal spike activity using Piphys. Spike train (black trace) from a recorded neuron in the time domain from Piphys. Spikes shown here are sorted from SpyKING CIRCUS software and labeled on the raw data with green and orange dots. Bottom: spike raster is aligned with the detected spikes showing firing activities at specific positions. (1)(2)(3) Individual spike examples randomly picked from the spike train.

To further demonstrate the applicability of Piphys to primary neuron recording, we compare the shape of the detected action potential and quality metrics such as amplitude distribution, interspike interval distribution, and firing rate to commercially available systems (Figure 6). The data was recorded from the same channel in the same well of neurons by Piphys, Intan, and Axion systems in sequential order on the same day. The data recorded on Piphys corresponds to the data obtained from both commercial systems, with high similarity to Intan and overall consistency with Axion across metrics in Figure 6.

**Figure 6.**
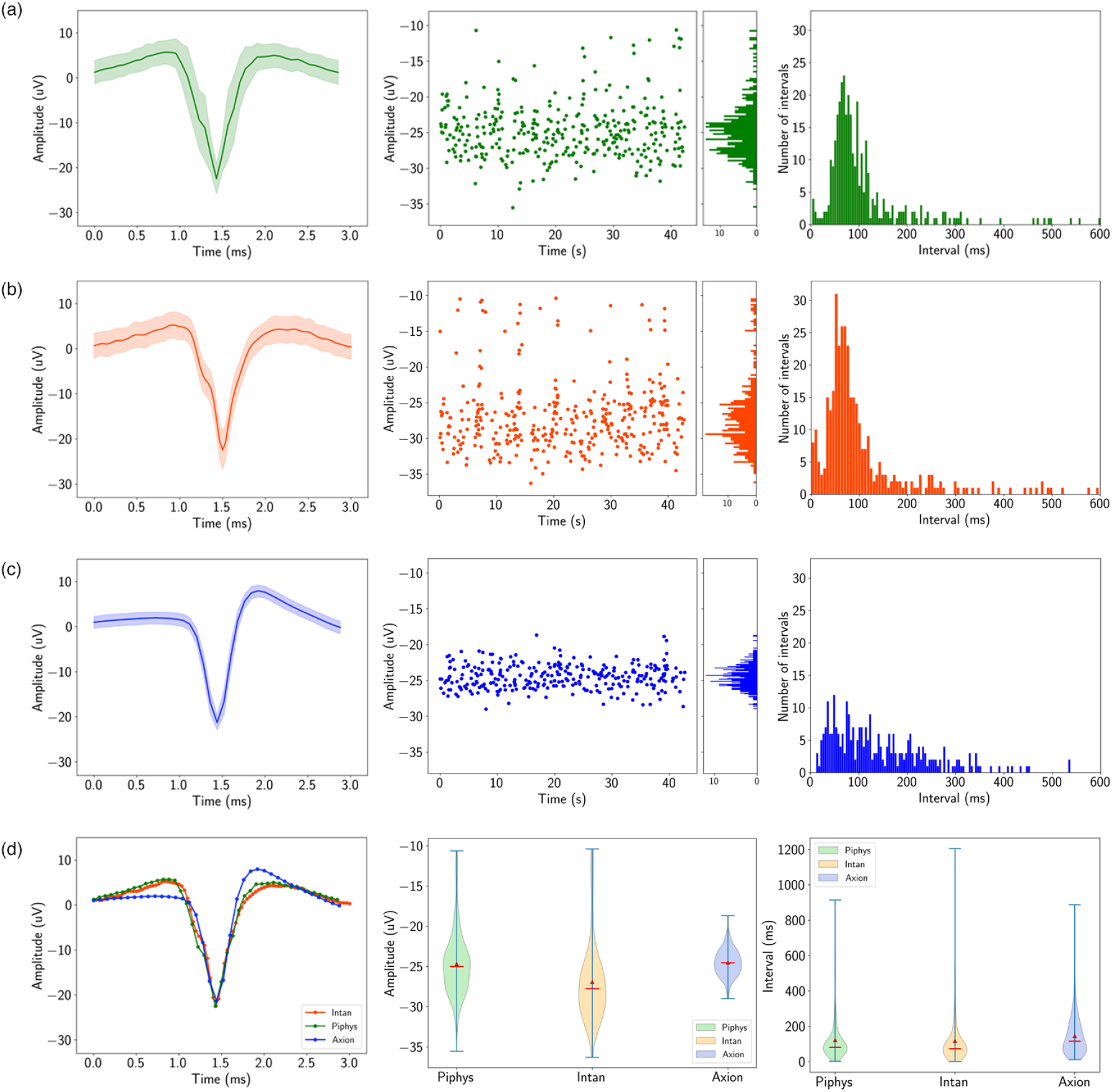
Piphys performance is similar to commercial systems. Spike sorting result for the same recording channel from Piphys, Intan RHD USB interface board, and Axion Maestro Edge. Shown from left to right are mean waveform with standard deviation (shaded area), amplitudes of the detected spikes over time, and interspike interval distribution. (a) Piphys (b) Intan RHD USB interface board (c) Axion Maestro Edge (d) Comparison of the mean waveform, amplitude, and interspike interval distribution from three systems.

The mean spike waveform, shown in the first column of Figure 6, was determined by averaging the voltage in a 3 ms window centered around the point where the voltage crossed the spike threshold. Differences in Axion’s waveform shape are a flatter starting point and a higher upstroke before settling to resting state. The amplitudes for the mean waveform are −24.67 ± 3.92 *μV* for Piphys, −26.92 ± 4.96 *μV* for Intan, and −24.50 ± 1.69 *μV* for Axion. Axion has a smaller deviation than Piphys and Intan, showing lower noise in the recording system.

The amplitudes of the detected spikes over time, shown in the middle column of Figure 6 are more sparse for Axion than for Intan and Piphys. Firing rates in events per second over the recording period shown are 8.05 for Piphys, 8.44 for Intan, and 6.86 for Axion.

The interspike interval histograms, shown in the middle column of Figure 6, have similar longer-tail distributions for Piphys and Intan centered at 122.79 ms and 118.15 ms, and a tighter distribution for Axion centered at 145.57 ms. However, the interspike interval means for all three systems are significantly close together.

The variation between Piphys and Axion could be attributed to physical differences in the circuity and possible advanced filtering performed by Axion’s proprietary BioCore v4 chip ‡. The filtering could account for the smoothness and low variability of the signal (measured 1.12 ± 0.18 *μV* RMS noise baseline), resulting in a smaller number of identified firing events with a tighter distribution. Piphys and Intan systems both use the same amplifier chips (Intan RHD2000 series), where the optional on-chip filtering was disabled during recording §. The raw signal, therefore, has a larger noise margin (measured 3.21 ± 0.66 *μ*V RMS noise baseline for Intan, 2.36 ± 0.4 *μ*V RMS for Piphys), which may create more false-positive firing events. The tail of the amplitude distributions in Intan and Piphys is skewed towards lower-amplitude events, closer to the noise floor. The interspike intervals for Intan and Piphys register several events with near-zero intervals, likely suggesting false-positive spikes from noise contamination. Contamination from noise, which is likely symmetrical, could affect the shape of the mean waveform calculated by overlaying and averaging all registered spikes.

Overall, these results demonstrate that Piphys can record neural activity in a manner comparable to commercially available hardware and software.

### Detecting burst activity from primary neuron network

On day 42 of culture, we recorded from the neurons with Piphys and found the primary neurons displayed synchronized network bursts, consistent with previous observations [35, 36]. Figure 7 shows the synchronous activity captured across four channels. After spike sorting, most detected spikes were arranged in short intervals with periods of silence in between. The spikes inside the bursts align among the channels, indicating that synchronized activity was present through the network. Quantitatively, the bursting has a general population rate of 0.13 bursts each second, with each burst lasting around 1 second. Within one burst, the number of spikes is 55 ± 17.58.

**Figure 7.**
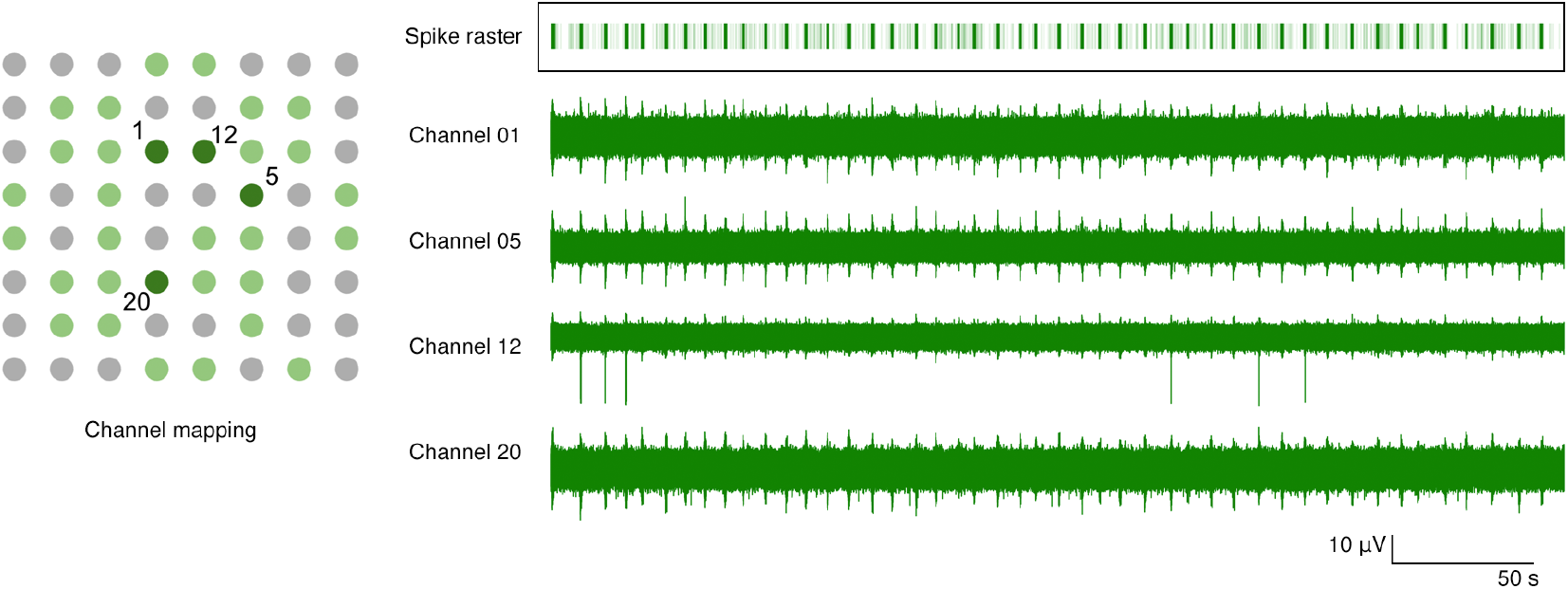
Bursting activity across four channels with channel mapping. Channel mapping shows 64 electrodes in well B2 of the Axion plate. Light green dots are the 32 electrodes recorded by Piphys. Dark green dots mark channels 1, 5, 12 and 20 whose raw recording plots are on the right. The spike raster superimposes all the detected spikes in the shown channels. Each light green vertical line in the raster indicates a spike, and the dark green bar is the result of superimposing multiple spikes in the burst. The bars in the raster plot align with the bursts throughout these four channels.

To further characterize Piphys system’s performance, we compute the SNR of bursting activity by the following equation applied to the smoothed signal:

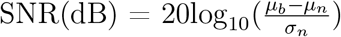

where *μ_b_* and *μ_n_* are the mean for the burst and baseline noise, respectively, *σ_n_* is the standard deviation of the noise. In Figure 8, background signal in green represents the original recording. The signal in blue is the smoothed product by boxcar averaging with a window size of three times the standard deviation of the original. The median SNR across active channels is measured at 4.35 dB. The mean for baseline noise in the burst recording is around 2.13 *μV* RMS, consistent with the noise measurement for the experiments described in the Performance section. These experiments further demonstrate that the Piphys system is sensitive and reliable in the relatively low amplitude neural signal recording range. In addition, with its open-source, light-weight, and remote monitoring capability through the IoT, Piphys adds unique value in extracellular electrophysiology.

**Figure 8.**
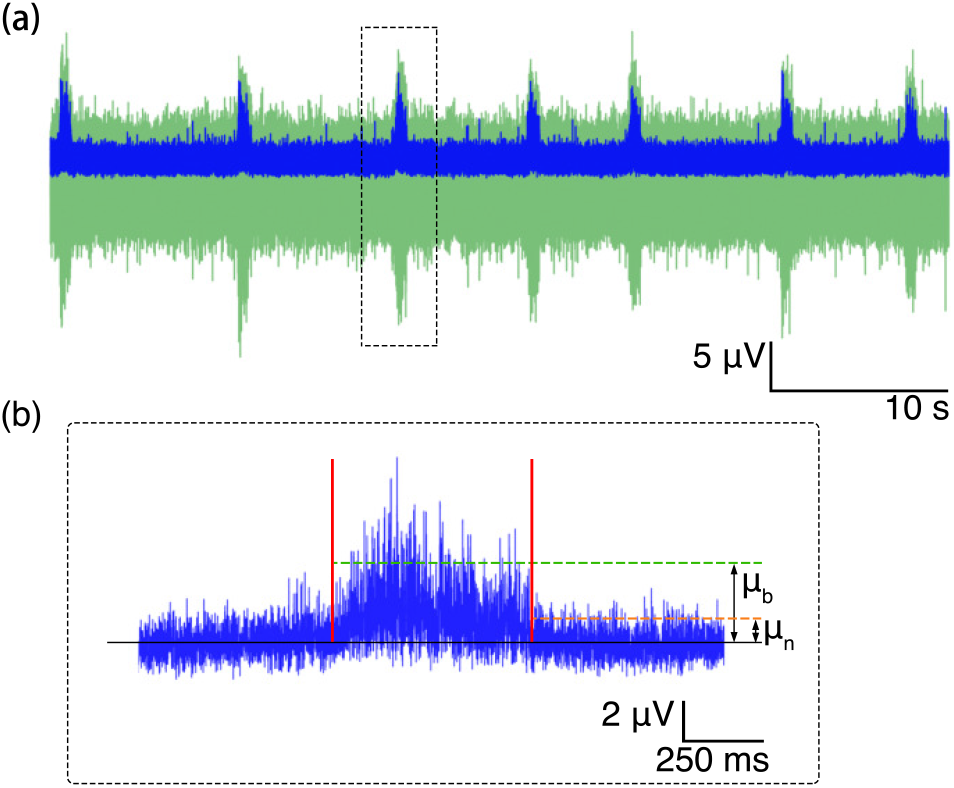
Signal-to-noise ratio of the burst. (a) Burst train (green) and the smoothed signal (blue). (b) Zoom in to the third smoothed burst showing means of the signal and the baseline noise for SNR calculation.

## 4. Comparison to other platforms

Comparing electrophysiology platforms side by side is challenging because each system fits a specific niche and requirements for a particular workflow. Different platforms arose as solutions to different problems, challenges, and user needs. Piphys arose due to the need for automation of experiments, integration with other IoT sensors, and flexible recording equipment that can be used in a fleet for longitudional study of many *in vitro* replicates.

Table 1 summarizes electrophysiology systems comparable to Piphys. The Axion Maestro Edge is designed as an out-of-the box bench top electrophysiology system with maximum comfort and usability. Although it has the highest price per channel, it is also an incubator. The Intan RHD USB interface board and headstages require more effort to calibrate, ground, and shield. Unlike Axion, Intan designs and code are open source. Intan bioamplifier chips have been used in many open source systems, including Intsy, Willow, Open Ephys, and now Piphys. Intsy was designed for measuring gastrointestinal (EGG), cardiac (ECG), neural (EEG), and neuromuscular (EMG) signals [17]. Willow was designed for high channel count neural probes and resolves the need for many computers by writing data directly to hard drives [20]. Open Ephys is an alternative system to Intan integrating more features into their GUI for closed-loop experiments and plugin-based workflows [18]^+^. Noise measurements for Piphys, Intan, and Axion were experimentally recorded, while noise measurements for Intsy, Willow, and Open Ephys were cited. Intan claims 2.4*μ*V RMS as typical in the datasheet for their chips * which was likely inherited into Open Ephys documentation. The whole system noise for Open Ephys is not explicitly mentioned in documentation.

**Table 1.** Comparison of Piphys features to several commercial and open source electrophysiology systems. Sampling Rate and Channels columns show the maximum numbers for all systems. * Noise shown on Open Ephys website is the amplifier input noise for Intan RHD2132 bioamplifier chip, not the whole system noise. ^†^RMS noise recorded experimentally.

| Platform | System Noise<br>($\mu\text{V RMS}$) | Sample Rate<br>(kHz) | Channels | Cost<br>(USD) | Cost<br>per<br>Channel | Open<br>Source | IoT |
| --- | --- | --- | --- | --- | --- | --- | --- |
| Pipphys | $2.36 \pm 0.4$ <sup>†</sup> | 15 | 32 | \$1,545 | \$48 | Yes | Yes |
| Intsy [17] | 6-8 | 2 | 64 | \$2,500 | \$39 | Yes | No |
| Intan RHD \|\| | $3.21 \pm 0.66$ <sup>†</sup> | 30 | 256 | \$10,295 | \$40 | Yes | No |
| Open Ephys [18] | 2.4 * | 30 | 512 | \$15,545 | \$30 | Yes | No |
| Willow [20] | 3.9 | 30 | 1024 | \$20,480 | \$20 | Yes | No |
| Axion Maestro Edge ¶ | $1.12 \pm 0.18$ <sup>†</sup> | 12.5 | 384 | \$70,000 | \$182 | No | No |

Piphys is the only electrophysiology device that supports Internet of Things (IoT) software integration out of the box. The IoT hardware modules and cloud software allow for horizontal scalability, enabling long-term observations of development, organization, and neural activity at scale, and integration with other IoT sensors. Piphys has a low entry cost, and the cost per channel can also be significantly lowered by increasing the number of channels supported per device. This would be accomplished by engineering an inexpensive FPGA into the controller shield to sample multiple bioamplifier chips and buffer those readings for the Pi. Piphys can have a large cost reduction if extra specialty connectors and adapters are removed (cutting roughly $300) and it is fitted with a less expensive USB cable.

## 5. Discussion

Remote longitudinal recording of neural circuits on an accessible platform will open up many exciting avenues for research into the physiology, organization, development, and adaptation of neural tissue. Integration with cloud software will allow in-depth experimentation and automation of analysis.

The proof of principle for Piphys has been shown on 2D cultures. As experiments with other devices have shown, it should be applicable to measurements of 3D brain organoids, which are becoming and increasingly popular model for studying human brain tissue development and function [5, 37–42].

Many electrode probes have been designed to interface with tissues to provide measurement points for voltage recordings [15, 33, 34, 43, 44]. Future work on Piphys would involve expanding the number of different electrodes types for long-term culture of the biological sample through collaborations with other research groups.

Future work on Piphys also includes increasing sampling rate and precision of timing in between samples. Currently, the Raspberry Pi CPU samples the Intan RHD2132 bioamplifier chip, and the sampling rates are limited by the CPU’s ability to multitask. Future solutions may involve adding another CPU or FPGA to the hardware shield. The platform will continue to be improved, and its modularity allows adapting hardware and software components as different needs arise.

Signal-to-noise ratio could be improved with enabling and tuning on-chip filtering, and improving Faraday cage shielding. *In vitro* cultures typically fire with amplitudes between 10 - 40 *μ*V [5, 7, 45]. They demand sensitive recording equipment, as an increase of just a few *μ*V in noise for spikes on the lower end of the spectrum can be considered a non-trivial variable.

Piphys software and hardware source files for building the Piphys system are available open source on GitHub ♯. All files are provided ‘as is’ and end-users are encouraged to freely use and adapt the system for their own application-specific protocols.

Overall, the open source Piphys design, programmability, and extreme flexibility of the Raspberry Pi significantly lowers the entry barrier of the electrophysiology system, providing an opportunity for broader applications in education and research.

## 6. Methods

### Tissue source

De-identified tissue samples were collected with previous patient consent in strict observance of the legal and institutional ethical regulations. Protocols were approved by the Human Gamete, Embryo, and Stem Cell Research Committee (institutional review board) at the University of California, San Francisco.

### Primary neuron culture

Prior to cell culture, the electrode surfaces of 6-well Axion plates (Axion Biosystems, CytoView MEA 6) were coated with 10 mg/mL poly-D-lysine (Sigma, P7280) at room temperature overnight. The following day, plates were rinsed 4x with water and dried at room temperature. Primary cells were obtained from human brain tissue at gestational week 21. Briefly, cortical tissue was cut into small pieces, incubated in 0.25% trypsin (Gibco, 25200056) for 30 minutes, then triturated in the presence of 10mg/mL DNAse (Sigma Aldrich, DN25) and passed through a 40um cell strainer. Cells were spun down and resuspended in BrainPhys (StemCell Technologies, 05790) supplemented with B27 (Thermo Fisher, 17504001), N2 (Thermo Fisher, 17502001), and penicillin-streptomycin (Thermo Fisher, 15070063), then diluted to a concentration of 8,000,000 cells/mL. Laminin (Thermo Fisher, 23017015, final concentration 50ug/mL) was added to the final aliquot of cells, and a 10uL drop of cells was carefully pipetted directly onto the dried, PDL-coated electrodes, forming an intact drop. The plate was transferred to a 37C, 5% CO2 incubator for 1 hour to allow the cells to settle, then 200uL of supplemented BrainPhys media was gently added to the drops. The following day, another 800uL of media was added, and each well was kept at 1 mL media for the duration of the cultures, with half the volume exchanged with fresh media every other day. Activity was first observed at 14 days in culture, and the second recordings were performed on day 42 of culture.

### Circuit board design, reduction of noise and EMI

The printed circuit board was designed in Autodesk Eagle. The board has four layers of copper. The top and bottom layers of the board are GND, while the two layers inside are signal and power. Every signal via has a ground via next to it to sink EMI as signals switch layers. The layout of the circuit board is done in modules. Via stitching was done around the perimeter and throughout the board area to separate modules (highlighted by squares in Figure 2) and fill in areas with no components. The amplifier chip and Raspberry Pi computer are separated by a cable such that noise from the computer would not interfere with the sensitive neural signal recording. During data acquisition, all of the electronics and biology were shielded by a 1 mm thick steel faraday cage.

### Cloud services integration

We deployed servers and cloud computing platforms to achieve permanent data storage and messaging between the local device and the dashboard. We used Remote Dictionary Server (Redis), Amazon Web Services Internet of Things (AWS IoT), and Simple Storage Service (S3). All services (except AWS IoT) are platform agnostic and can be hosted anywhere. For our particular experimental setup, Redis and S3 were hosted on the Pacific Research Platform (PRP) [28]. The Internet of Things (IoT) service with MQTT messaging and device management was coordinated through Amazon Web Services (AWS). The dashboard was hosted on a server at UC Santa Cruz.

### Redis, real-time data stream

Neuronal action potential recording with a high sample rate and multiple channels requires a high throughput pipeline to make realtime streaming possible. Remote Dictionary Server (Redis) is a good choice for the implementation of this objective. It is a high-speed cloud-based data structure store that can be used as a cache, message broker, and database. Based on benchmarking results, Redis can handle hundreds of thousands of requests per second. The highest data rate for every push from Piphys system to Redis is 9.6 MB (32 channels × 15 kHz sampling rate × 16 bits/sample × 10 seconds), which can be satisfied with an internet bandwidth larger than 7.68 Mbps.

### Internet of Things (IoT) communication

The dashboard is programmed to be an IoT device that sends Message Queuing Telemetry Transport (MQTT) messages to control and check the Piphys system. In response, the Piphys subscribes to a particular MQTT topic to wait for instructions. The AWS IoT supports the communication of hundreds of devices, making the Piphys system’s extension to a large scale possible in the future.

### Simple Storage Service (S3)

The Simple Storage Service (S3) is the final data storage location. S3 is accessible from anywhere at any time on the internet. It supports both management from a terminal session and integration to a custom web browser application. After each experiment, a new identifier will be updated on the dashboard. When a user asks for a specific experiment result, the dashboard can pull the corresponding data file directly from S3 for visualization.

## Acknowledgements

This work is supported by the Schmidt Futures Foundation SF 857 (D.H.). Research reported in this publication was also supported by the National Institute Of Mental Health of the National Institutes of Health under award number R01MH120295 (S.R.S.), the National Science Foundation under award number NSF 2034037 (M.T.), the National Defense Science and Engineering Graduate Fellowship (00002116, M.G.K.), and gifts from Schmidt Futures and the William K. Bowes Jr Foundation (T.J.N). K.V. was supported by grant T32HG008345 from the National Human Genome Research Institute (NHGRI), part of National Institutes of Health (NIH), USA. D.H. is a Howard Hughes Medical Institute Investigator.

We are thankful to the Pacific Research Platform, supported by the National Science Foundation under award number NSF 1541349.

We would like to thank Rob Currie for contributing early advice and inspiration, Erik Jung for encouragement and organizational support, Dr. Alex Pollen and Dr. Mohammed A. Mostajo-Radji for sharing their Axion system, and Pattawong Pansodtee for help with CNC fabrication of the plate holder.

## Author contributions statement

K.V. created the electronics designs, J.G. wrote the device software, M.G.K. cultured the primary neurons, D.F.P. and S.E.S. created the streaming dashboard. J.G., K.V, and M.G.K. ran the experiments. J.G.and N.H. analysed the results. D.B.F. advised electronics design, and M.T. advised the mechanical design. D.H., M.T., S.R.S, and T.J.N. supervised the team and secured funding. K.V., J.G., and M.G.K. wrote the manuscript with support from D.F.P, T.J.N, and D.H. All authors reviewed the manuscript.

## Footnotes

‡ https://www.axionbiosystems.com/resources/product-brochure/maestro-edge-mea-system-brochure

§ https://intantech.com/files/Intan_RHD2000_series_datasheet.pdf

+ https://open-ephys.org

* https://intantech.com/files/Intan_RHD2000_series_datasheet.pdf

♯ https://github.com/braingeneers/piphys

## References

[1] D. Hansel, G. Mato, and C. Meunier. Synchrony in Excitatory Neural Networks. Neural Computation, 7(2):307–337, March 1995. Publisher: MIT Press.

[2] John M. Beggs and Dietmar Plenz. Neuronal Avalanches in Neocortical Circuits. Journal of Neuroscience, 23(35):11167–11177, December 2003. Publisher: Society for Neuroscience Section: Behavioral/Systems/Cognitive.

[3] Daniele Poli, Vito P. Pastore, and Paolo Massobrio. Functional connectivity in in vitro neuronal assemblies. Frontiers in Neural Circuits, 9, 2015. Publisher: Frontiers.

[4] Yu-Ting Huang, Yu-Lin Chang, Chun-Chung Chen, Pik-Yin Lai, and C. K. Chan. Positive feedback and synchronized bursts in neuronal cultures. PLOS ONE, 12(11):e0187276, November 2017. Publisher: Public Library of Science.

[5] Cleber A. Trujillo, Richard Gao, Priscilla D. Negraes, Jing Gu, Justin Buchanan, Sebastian Preissl, Allen Wang, Wei Wu, Gabriel G. Haddad, Isaac A. Chaim, Alain Domissy, Matthieu Vandenberghe, Anna Devor, Gene W. Yeo, Bradley Voytek, and Alysson R. Muotri. Complex Oscillatory Waves Emerging from Cortical Organoids Model Early Human Brain Network Development. Cell Stem Cell, 25(4):558–569.e7, October 2019.

[6] Tal Sharf, Tjitse van der Molen, Elmer Guzman, Stella M. K. Glasauer, Gabriel Luna, Zhouwei Cheng, Morgane Audouard, Kamalini G. Ranasinghe, Kiwamu Kudo, Srikantan S. Nagarajan, Kenneth R. Tovar, Linda R. Petzold, Paul K. Hansma, and Kenneth S. Kosik. Intrinsic network activity in human brain organoids. bioRxiv, page 2021.01.28.428643, January 2021. Publisher: Cold Spring Harbor Laboratory Section: New Results.

[7] Joseph Negri, Vilas Menon, and Tracy L. Young-Pearse. Assessment of Spontaneous Neuronal Activity In Vitro Using Multi-Well Multi-Electrode Arrays: Implications for Assay Development. eNeuro, 7(1), January 2020.

[8] Yuichiro Yada, Takeshi Mita, Akihiro Sanada, Ryuichi Yano, Ryohei Kanzaki, Douglas J. Bakkum, Andreas Hierlemann, and Hirokazu Takahashi. Development of neural population activity toward self-organized criticality. Neuroscience, 343:55–65, 2017.

[9] Maria-Patapia Zafeiriou, Guobin Bao, James Hudson, Rashi Halder, Alica Blenkle, Marie-Kristin Schreiber, Andre Fischer, Detlev Schild, and Wolfram-Hubertus Zimmermann. Developmental GABA polarity switch and neuronal plasticity in Bioengineered Neuronal Organoids. Nature Communications, 11(1):3791, July 2020. Number: 1 Publisher: Nature Publishing Group.

[10] Friedemann Zenke and Wulfram Gerstner. Hebbian plasticity requires compensatory processes on multiple timescales. Philosophical Transactions of the Royal Society of London. Series B, Biological Sciences, 372(1715), 2017.

[11] Ashesh K Dhawale, Rajesh Poddar, Steffen BE Wolff, Valentin A Normand, Evi Kopelowitz, and Bence P Ölveczky. Automated long-term recording and analysis of neural activity in behaving animals. eLife, 6:e27702, September 2017. Publisher: eLife Sciences Publications, Ltd.

[12] Maithra Raghu and Eric Schmidt. A Survey of Deep Learning for Scientific Discovery. arXiv:2003.11755 [cs, stat], March 2020. arXiv: 2003.11755.

[13] Konstantinos Nasiotis, Martin Cousineau, François Tadel, Adrien Peyrache, Richard M. Leahy, Christopher C. Pack, and Sylvain Baillet. Integrated opensource software for multiscale electrophysiology. Scientific Data, 6(1):231, October 2019. Number: 1 Publisher: Nature Publishing Group.

[14] Joshua H Siegle, Gregory J Hale, Jonathan P Newman, and Jakob Voigts. Neural ensemble communities: open-source approaches to hardware for large-scale electrophysiology. Current Opinion in Neurobiology, 32:53–59, June 2015.

[15] Jan Putzeys, Bogdan C. Raducanu, Alain Carton, Jef De Ceulaer, Bill Karsh, Joshua H. Siegle, Nick Van Helleputte, Timothy D. Harris, Barundeb Dutta, Silke Musa, and Carolina Mora Lopez. Neuropixels Data-Acquisition System: A Scalable Platform for Parallel Recording of 10 000+ Electrophysiological Signals. IEEE Transactions on Biomedical Circuits and Systems, 13(6):1635–1644, December 2019.

[16] Taiga Abe, Ian Kinsella, Shreya Saxena, Liam Paninski, and John P Cunningham. Neuroscience Cloud Analysis As a Service. page 32.

[17] Jonathan C. Erickson, James A. Hayes, Mauricio Bustamante, Rajwol Joshi, Alfred Rwagaju, Niranchan Paskaranandavadivel, and Timothy R. Angeli. Intsy: a low-cost, open-source, wireless multi-channel bioamplifier system. Physiological Measurement, 39(3):035008, March 2018. Publisher: IOP Publishing.

[18] Joshua H Siegle, Aarón Cuevas López, Yogi A Patel, Kirill Abramov, Shay Ohayon, and Jakob Voigts. Open Ephys: an open-source, plugin-based platform for multichannel electrophysiology. Journal of Neural Engineering, 14(4):045003, August 2017.

[19] Jonathan Paul Newman, Riley Zeller-Townson, Ming-fai Fong, Sharanya Arcot Desai, Robert E. Gross, and Steve M. Potter. Closed-Loop, Multichannel Experimentation Using the Open-Source NeuroRighter Electrophysiology Platform. Frontiers in Neural Circuits, 6, 2013. Publisher: Frontiers.

[20] Justin P. Kinney, Jacob G. Bernstein, Andrew J. Meyer, Jessica B. Barber, Marti Bolivar, Bryan Newbold, Jorg Scholvin, Caroline Moore-Kochlacs, Christian T. Wentz, Nancy J. Kopell, and Edward S. Boyden. A direct-to-drive neural data acquisition system. Frontiers in Neural Circuits, 9, 2015.

[21] Leonardo D. Garma, Laura Matino, Giovanni Melle, Fabio Moia, Francesco De Angelis, Francesca Santoro, and Michele Dipalo. Cost-effective and multifunctional acquisition system for in vitro electrophysiological investigations with multielectrode arrays. PLOS ONE, 14(3):e0214017, March 2019. Publisher: Public Library of Science.

[22] A. F. Hussein, N. A. kumar, M. Burbano-Fernandez, G. Ramírez-González, E. Abdulhay, and V. H. C. De Albuquerque. An Automated Remote Cloud-Based Heart Rate Variability Monitoring System. IEEE Access, 6:77055–77064, 2018. Conference Name: IEEE Access.

[23] Zhe Yang, Qihao Zhou, Lei Lei, Kan Zheng, and Wei Xiang. An IoT-cloud Based Wearable ECG Monitoring System for Smart Healthcare. Journal of Medical Systems, 40(12):286, October 2016.

[24] M. Hassanalieragh, A. Page, T. Soyata, G. Sharma, M. Aktas, G. Mateos, B. Kantarci, and S. Andreescu. Health Monitoring and Management Using Internet-of-Things (IoT) Sensing with Cloud-Based Processing: Opportunities and Challenges. In 2015 IEEE International Conference on Services Computing, pages 285–292, June 2015.

[25] A. A. P. Wai, H. Dajiang, and N. S. Huat. IoT-enabled multimodal sensing headwear system. In 2018 IEEE 4th World Forum on Internet of Things (WF-IoT), pages 286–290, February 2018.

[26] J. Hermiz, N. Rogers, E. Kaestner, M. Ganji, D. Cleary, J. Snider, D. Barba, S. Dayeh, E. Halgren, and V. Gilja. A clinic compatible, open source electrophysiology system. In 2016 38th Annual International Conference of the IEEE Engineering in Medicine and Biology Society (EMBC), pages 4511–4514, August 2016. ISSN: 1558-4615.

[27] P. K. Yong and E. T. Wei Ho. Streaming brain and physiological signal acquisition system for IoT neuroscience application. In 2016 IEEE EMBS Conference on Biomedical Engineering and Sciences (IECBES), pages 752–757, December 2016.

[28] Larry Smarr, Camille Crittenden, Thomas DeFanti, John Graham, Dmitry Mishin, Richard Moore, Philip Papadopoulos, and Frank Würthwein. The Pacific Research Platform: Making High-Speed Networking a Reality for the Scientist. In Proceedings of the Practice and Experience on Advanced Research Computing, PEARC ‘18, pages 1–8, Pittsburgh, PA, USA, July 2018. Association for Computing Machinery.

[29] D. Wagenaar, T.B. DeMarse, and S.M. Potter. MeaBench: A toolset for multielectrode data acquisition and on-line analysis. In Conference Proceedings. 2nd International IEEE EMBS Conference on Neural Engineering, 2005., pages 518–521, March 2005. ISSN: 1948-3554.

[30] Jason E. Chung, Jeremy F. Magland, Alex H. Barnett, Vanessa M. Tolosa, Angela C. Tooker, Kye Y. Lee, Kedar G. Shah, Sarah H. Felix, Loren M. Frank, and Leslie F. Greengard. A Fully Automated Approach to Spike Sorting. Neuron, 95(6):1381–1394.e6, September 2017.

[31] Pierre Yger, Giulia LB Spampinato, Elric Esposito, Baptiste Lefebvre, Stéphane Deny, Christophe Gardella, Marcel Stimberg, Florian Jetter, Guenther Zeck, Serge Picaud, Jens Duebel, and Olivier Marre. A spike sorting toolbox for up to thousands of electrodes validated with ground truth recordings in vitro and in vivo. eLife, 7:e34518, March 2018. Publisher: eLife Sciences Publications, Ltd.

[32] JinHyung Lee, Catalin Mitelut, Hooshmand Shokri, Ian Kinsella, Nishchal Dethe, Shenghao Wu, Kevin Li, Eduardo B. Reyes, Denis Turcu, Eleanor Batty, Young J. Kim, Nora Brackbill, Alexandra Kling, Georges Goetz, E. J. Chichilnisky, David Carlson, and Liam Paninski. YASS: Yet Another Spike Sorter applied to large-scale multi-electrode array recordings in primate retina. bioRxiv, page 2020.03.18.997924, March 2020. Publisher: Cold Spring Harbor Laboratory Section: New Results.

[33] Yoonseok Park, Colin K. Franz, Hanjun Ryu, Haiwen Luan, Kristen Y. Cotton, Jong Uk Kim, Ted S. Chung, Shiwei Zhao, Abraham Vazquez-Guardado, Da Som Yang, Kan Li, Raudel Avila, Jack K. Phillips, Maria J. Quezada, Hokyung Jang, Sung Soo Kwak, Sang Min Won, Kyeongha Kwon, Hyoyoung Jeong, Amay J. Bandodkar, Mengdi Han, Hangbo Zhao, Gabrielle R. Osher, Heling Wang, KunHyuck Lee, Yihui Zhang, Yonggang Huang, John D. Finan, and John A. Rogers. Three-dimensional, multifunctional neural interfaces for cortical spheroids and engineered assembloids. Science Advances, 7(12):eabf9153, March 2021. Publisher: American Association for the Advancement of Science Section: Research Article.

[34] Long Yang, Kwang Lee, Jomar Villagracia, and Sotiris C. Masmanidis. Open source silicon microprobes for high throughput neural recording. Journal of Neural Engineering, 17(1):016036, January 2020. Publisher: IOP Publishing.

[35] Daniel A. Wagenaar, Radhika Madhavan, Jerome Pine, and Steve M. Potter. Controlling Bursting in Cortical Cultures with Closed-Loop Multi-Electrode Stimulation. The Journal of Neuroscience, 25(3):680–688, January 2005.

[36] Douglas J. Bakkum, Milos Radivojevic, Urs Frey, Felix Franke, Andreas Hierlemann, and Hirokazu Takahashi. Parameters for burst detection. Frontiers in Computational Neuroscience, 7, 2014. Publisher: Frontiers.

[37] Mototsugu Eiraku, Kiichi Watanabe, Mami Matsuo-Takasaki, Masako Kawada, Shigenobu Yonemura, Michiru Matsumura, Takafumi Wataya, Ayaka Nishiyama, Keiko Muguruma, and Yoshiki Sasai. Self-Organized Formation of Polarized Cortical Tissues from ESCs and Its Active Manipulation by Extrinsic Signals. Cell Stem Cell, 3(5):519–532, November 2008. Publisher: Elsevier.

[38] Madeline A. Lancaster, Magdalena Renner, Carol-Anne Martin, Daniel Wenzel, Louise S. Bicknell, Matthew E. Hurles, Tessa Homfray, Josef M. Penninger, Andrew P. Jackson, and Juergen A. Knoblich. Cerebral organoids model human brain development and microcephaly. Nature, 501(7467):373–379, September 2013. Number: 7467 Publisher: Nature Publishing Group.

[39] Giorgia Quadrato, Tuan Nguyen, Evan Z. Macosko, John L. Sherwood, Sung Min Yang, Daniel Berger, Natalie Maria, Jorg Scholvin, Melissa Goldman, Justin Kinney, Edward S. Boyden, Jeff Lichtman, Ziv M. Williams, Steven A. McCarroll, and Paola Arlotta. Cell diversity and network dynamics in photosensitive human brain organoids. Nature, 545(7652):48–53, May 2017.

[40] Alex Shcheglovitov, Yueqi Wang, Laura Bell, Chad Russell, Celeste Armstrong, and Jay Spampanato. Human cortical organoids from single iPSC-derived neural rosettes for studying human cortical development and disorders. The FASEB Journal, 33(1_supplement):205.3–205.3, April 2019. Publisher: Federation of American Societies for Experimental Biology.

[41] Hideya Sakaguchi, Yuki Ozaki, Tomoka Ashida, Takayoshi Matsubara, Naotaka Oishi, Shunsuke Kihara, and Jun Takahashi. Self-Organized Synchronous Calcium Transients in a Cultured Human Neural Network Derived from Cerebral Organoids. Stem Cell Reports, 13(3):458–473, June 2019.

[42] Abraam M. Yakoub. Cerebral organoids exhibit mature neurons and astrocytes and recapitulate electrophysiological activity of the human brain. Neural Regeneration Research, 14(5):757, May 2019. Publisher: Wolters Kluwer – Medknow Publications.

[43] Qiang Li, Kewang Nan, Paul Le Floch, Zuwan Lin, Hao Sheng, and Jia Liu. Cyborg Organoids: Implantation of Nanoelectronics via Organogenesis for Tissue-Wide Electrophysiology. bioRxiv, page 697664, July 2019. Publisher: Cold Spring Harbor Laboratory Section: New Results.

[44] Jung Min Lee, Guosong Hong, Dingchang Lin, Thomas G. Schuhmann, Andrew T. Sullivan, Robert D. Viveros, Hong-Gyu Park, and Charles M. Lieber. Nanoenabled Direct Contact Interfacing of Syringe-Injectable Mesh Electronics. Nano Letters, 19(8):5818–5826, August 2019.

[45] Galina Schmunk, Chang N. Kim, Sarah S. Soliman, Matthew G. Keefe, Derek Bogdanoff, Dario Tejera, Ryan S. Ziffra, David Shin, Denise E. Allen, Bryant B. Chhun, Christopher S. McGinnis, Ethan A. Winkler, Adib A. Abla, Edward F. Chang, Zev J. Gartner, Shalin B. Mehta, Xianhua Piao, Keith B. Hengen, and Tomasz J. Nowakowski. Human microglia upregulate cytokine signatures and accelerate maturation of neural networks. bioRxiv, page 2020.03.24.006874, March 2020. Publisher: Cold Spring Harbor Laboratory Section: New Results.

